# Targeted, collaborative biodiversity conservation in the global ocean can benefit fisheries economies

**DOI:** 10.1101/2021.04.23.441004

**Authors:** D. Scott Rinnan, Gabriel Reygondeau, Jennifer McGowan, Vicky Lam, Rashid Sumaila, Ajay Ranipeta, Kristin Kaschner, Cristina Garilao, William L. Cheung, Walter Jetz

**Affiliations:** Department of Ecology and Evolutionary, Yale University, New Haven, CT, USA; Center for Biodiversity and Global Change, Yale University, New Haven, CT, USA; Institute for the Oceans and Fisheries, Changing Ocean Research Unit, University of British Columbia, Vancouver, British Columbia, Canada; The Nature Conservancy, Arlington, VA, USA; Institute for the Oceans and Fisheries & the School of Public Policy and Global Affairs, Fisheries Economics Research Unit, University of British Columbia, Vancouver, British Columbia, Canada; Department of Biometry & Environmental System Analysis, Albert Ludwigs University, Freiburg, Freiburg, Germany; GEOMAR Helmholtz Centre for Ocean Research Kiel, Kiel, Germany

## Abstract

Marine protected areas (MPAs) are key to averting continued loss of species and ecosystem services in our oceans, but concerns around economic trade-offs hamper progress. Here we provide optimized planning scenarios for global MPA networks that secure species habitat while minimizing impacts on fisheries revenues. We found that MPA coverage requirements differ vastly among nations, and that two-thirds of nations benefit economically from a collaborative approach. Immediate global protection of marine biodiversity habitat comes with losses of ~19% of total fisheries revenues, but international cooperation in concert with high seas protection improves economic losses for most countries, safeguards all species, and could save ~5B USD annually worldwide. Nations and fishery economies both share benefits from a coordinated approach to conserving marine biodiversity, with direct relevance to current international policies.

## Main Text

Nearly 40 years after the United Nations Convention on the Law of the Seas (UNCLOS) established the global ocean governance system, marine biodiversity is experiencing immense anthropogenic pressures (*1–4*). Fishing, climate change, pollution, coastal development, extractive industries and other human activities have been driving shifts in species ranges, changes in growth and reproduction (*5*), alteration of biological communities (*6, 7*), as well as loss of biomass (*8*), and ecosystem functions and services (*3, 9*).

Marine protected areas (MPAs) are a key scalable management mechanism to mitigate these human impacts (*10*), but coverage of the current MPA network is small and sporadic (<8% of the global ocean (*11*)), and varies dramatically within national waters, resulting in inadequate (*12*) and inefficient biodiversity representation (*13*). These shortcomings have led to calls for ambitious conservation targets, such as the Half-Earth Project’s recommendation that 50% of land and sea be under some form of long-term protection (*14*), and more recently a 30% protection target by 2030 (*15, 16*) now provisionally included under the Convention on Biological Diversity’s post-2020 Global Biodiversity Framework (*17*). The spatial conservation planning community has responded by highlighting conservation gaps and opportunity areas for marine species (*12, 18, 19*), and by performing spatial optimization to identify biodiversity priorities (*15, 19–21*) and multi-objective optimization for food security and carbon sequestration (*22*).

Yet for an international MPA network to be operationalizable and effective, we must also provide assessments of trade-offs with alternative uses (*16*). Fishing industries provide direct means of livelihood to an estimated 59.5 million people — producing upwards of 158 million tonnes of fish for global consumption in 2018 (*23*) — and contribute 192B USD to the global economy annually^1^ (*24*). Addressing the opportunity costs of foregoing profit from fishing is critical for framing MPAs as an economically viable and sustainable ocean conservation strategy (*16*), and a joint assessment of effectiveness and costs of future MPAs is urgently needed to support national policy commitments.

Here we identify these trade-offs and resulting opportunities for MPA expansion. We address key questions about jurisdictional responsibility, placement, and the economic impacts of marine protections on fisheries. We developed a spatial optimization framework to identify biodiversity priority areas (those that sufficiently represent marine mammals and fishes) for three different strategies: by adding additional protection to i) areas only within national jurisdictions (“Exclusive Economic Zones”, “EEZs”); ii) areas only beyond national jurisdiction (“high seas”); and iii) globally including both national waters and the high seas (“global ocean”). We optimized each strategy with two distinct socioeconomic objectives: minimizing costs related to area (*MinArea*) and minimizing the annual revenue loss to fisheries (*MinCost*, averaged 2007– 2016), under the baseline assumption that fishing pressure is removed with area-based conservation. We used the most up-to-date and comprehensive fisheries catch data (*25*) coupled with global marine ex-vessel fish price datasets (*26*) to quantify economic cost, and used existing MPAs (*11*) and anthropogenic impacts on marine ecosystems (*1*) in our assessments (Fig. S1).

Under the EEZs and global-ocean strategies we found large gains in species coverage could be made with small additions to existing MPAs, regardless of the conservation objective. For example, the number of current species meeting area-based representation targets could be doubled by protecting just an additional 3% of strategically-placed ocean area (Fig. 1A). In contrast, as the high seas has lower species richness and more wide-ranging species, more area is needed to meet species’ targets. Specifically, the high-seas strategy was only able to sufficiently protect 46% of all study species and 50% of species found in the high seas under either objective, largely via prioritization of high sea biodiversity hotspots (e.g., seamounts) that border EEZs (Fig. 2B, Fig. S2) and are already recognized for their conservation importance (*27*).

**Fig. 1.**
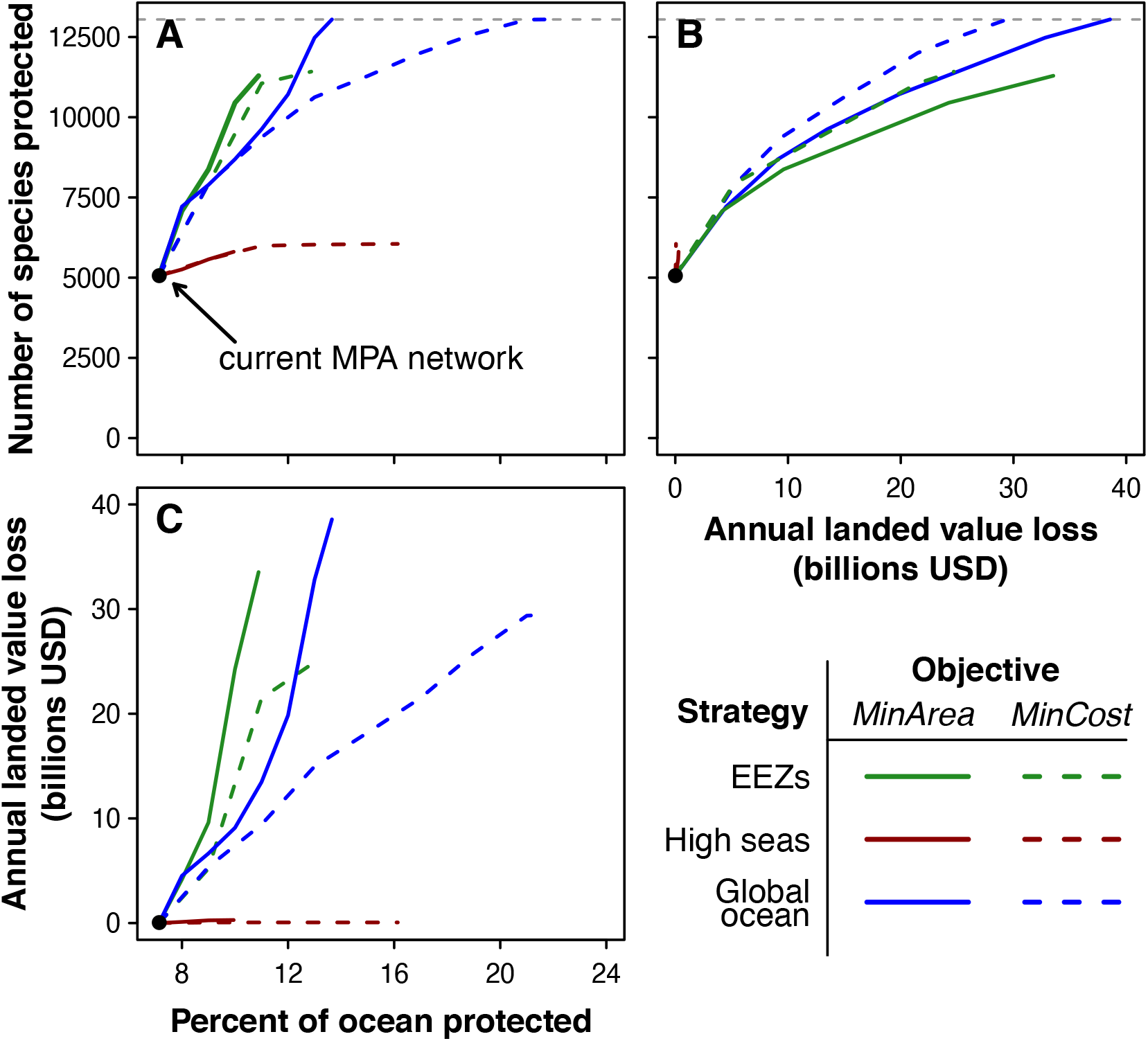
Accumulation curves illustrating the most efficient trajectory to optimality for each conservation strategy and cost objective. (A) The number of species with a sufficient amount of protected habitat as MPAs are added by priority rank. The grey dashed line represents all study species (n = 13046). (B) Number of species protected vs. loss in annual landed value of fisheries (averaged 2007–2016). (C) Loss in landed value of fisheries as MPAs are added.

**Fig. 2.**
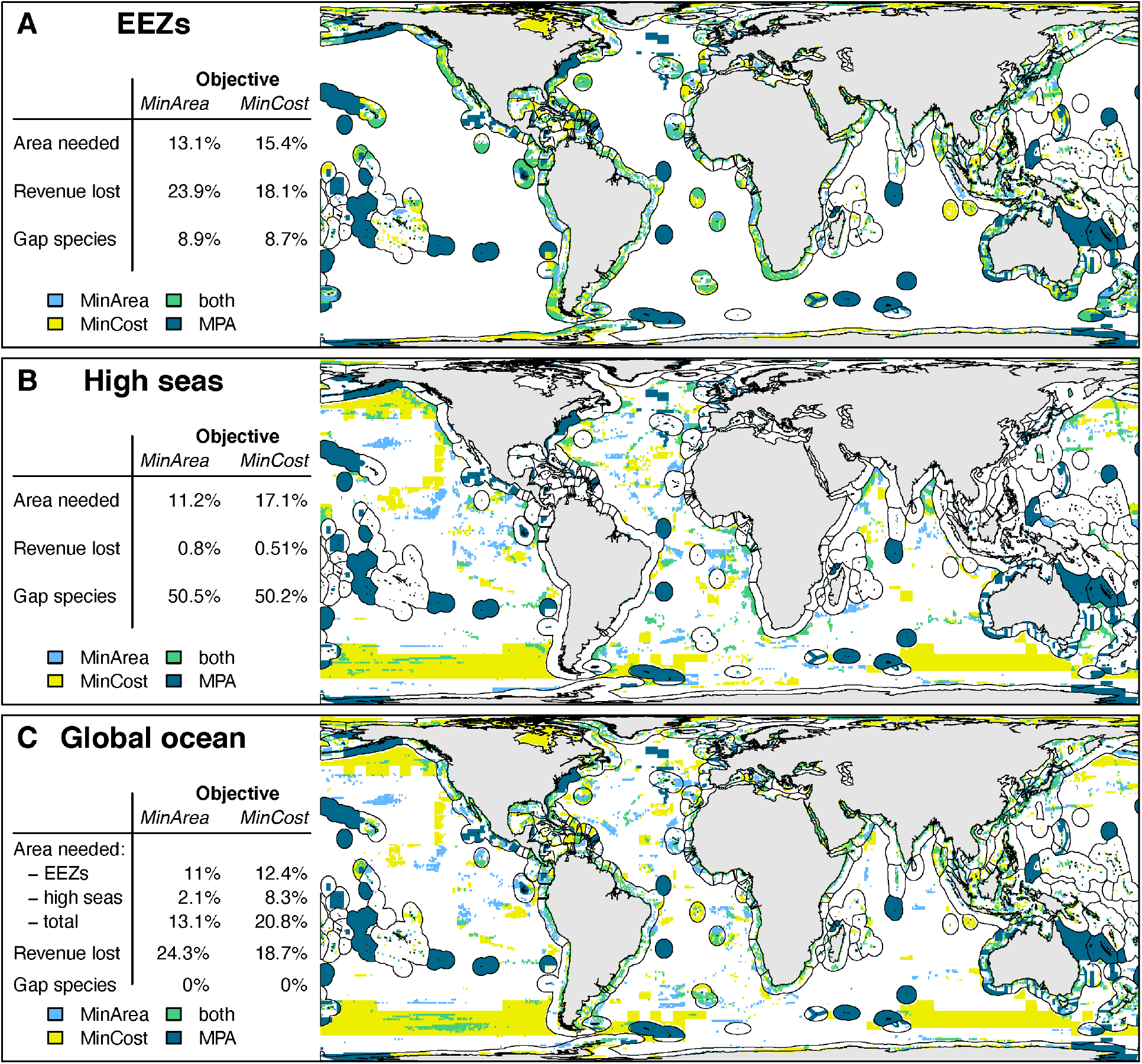
Priority conservation areas for three conservation strategies, focused on (A) areas only within national jurisdictions, (B) areas only beyond national jurisdiction, and (C) globally, including both national waters and the high seas. Each strategy identifies priority areas for species representation across two conservation objectives: minimizing area or revenue loss (cost) to fisheries. Areas selected with both objectives shown in green and current MPAs in dark blue.

The *MinArea* objective always resulted in more spatially efficient results. Under the EEZs strategy 13.1% of the ocean was required to meet the conservation targets for 91.1% of study species, with an expected annual 33.5B USD loss in revenue to the fishing industry (Fig. 1B,C). The global-ocean strategy closed the species protection gap, meeting 100% species representation over slightly less area, but resulted in an additional $5B expected revenue loss.

The *MinCost* objective, by contrast, required more area to meet the same levels of species representation regardless of the strategy (Fig. 1A; Fig. 2A). After rapid early gains offered by initial high-priority locations, minimizing loss of fisheries revenue protected the same number of species while preventing on average $9 billion in lost annual revenue compared to the *MinArea* objective for both the EEZs-only and all-oceans strategies. Regardless of objective, the high seas-only strategy resulted in marginal revenue loss to fisheries (<1%) (Fig. 1C) but secured minimal conservation gains beyond the current MPA network.

These comparisons between objectives offer at least two critical insights. First, strategic protection of the ocean is not inherently incompatible with the socioeconomic objectives of fisheries. The safeguarding of marine species could be achieved through strategic protection of 20.8% of the global ocean — of which 7.1% is already established or proposed as protected — while retaining 81.3% of current global fisheries revenues. This area is distributed across 12.4% of EEZ waters and 8.3% of the high seas, a large portion of which is relatively unexploited marine mammal habitat in the Southern Ocean and areas with lower human impacts such as the Arctic Ocean and Bering Sea (Fig. 2C), and these numbers are consistent with other prioritization results (*22*). Second, it is only through a coordinated global-ocean strategy that biodiversity can be sufficiently safeguarded. Due to the partial restriction of species to pelagic habitats (*28*), a conservation strategy focused solely on EEZ protection is at minimum inefficient and, for purely pelagic species, ineffective. This fact reinforces the crucial need of common conservation strategies between neighboring countries that share common stocks and non-exploited species (*29*).

Assessing further the representation of species in the *MinCost* solution we identify striking differences across the three strategies (Fig. 3). Most high-value commercial species can be sufficiently protected within EEZs, while many noncommercial species will be missed entirely. By contrast, limiting conservation to the high seas results in insufficient protection of the majority of both commercial and noncommercial species (Fig. 3), jeopardizing the viability of important fished species, for example the anchoveta (*Engraulis ringens*). For some species with small ranges that span national boundaries — such as the Gulf menhaden (*Brevoortia patronus*) — a global-ocean strategy is the only path toward successful conservation. Thus, sufficient representation of all marine species requires a coordinated global-ocean conservation strategy.

**Fig. 3.**
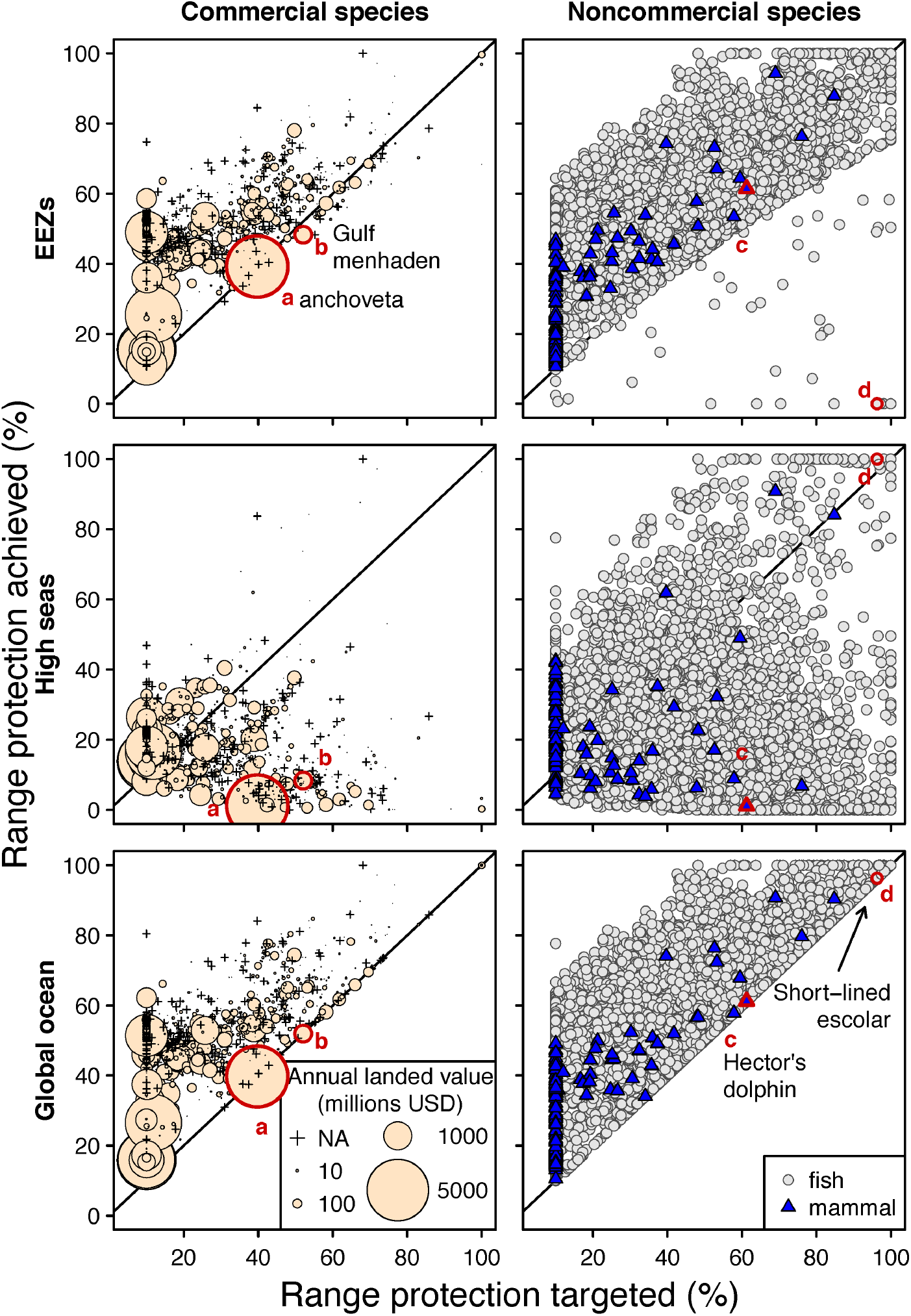
Conservation outcomes for commercial and noncommercial species under each conservation strategy (*MinCost* objective). The comparison relates spatial representation achieved (expressed in percent of habitat area conserved) to that targeted; species above the 1:1 line are those for which the conservation strategy achieved their representation targets.

Commercial species are shown with point size scaled by total landed value in millions of USD. Example species outlined in red: (a) *Engraulis ringens*, (b) *Brevoortia patronus*, (c) *Cephalorhynchus hectori*, (d) *Rexea brevilineata*.

For nations making commitments toward an expanded MPA network, our results offer insight around required individual contributions and how responsibilities may be affected by shared resources — such as invoking the extra-jurisdictional high seas. We found that the EEZs strategy required that countries on average protect an additional 22.6% of their own area (SD = 20.3%) to ensure full species representation, compared to 20.1% (SD = 19.5%) if a coordinated global-ocean strategy were embraced and operationalized (Fig. 4). This area-based efficiency benefited 37% of nations, with noteworthy examples including South Africa (ZAF), Ecuador (ECU), and Costa Rica (CRI). For the 47% of countries in which the global-ocean strategy necessitates greater areal protection commitments, these increases were small (mean = 3.2%, SD = 5.7%), but notable examples include Belize (BLZ), Romania (ROU), and Estonia (EST).

**Fig. 4.**
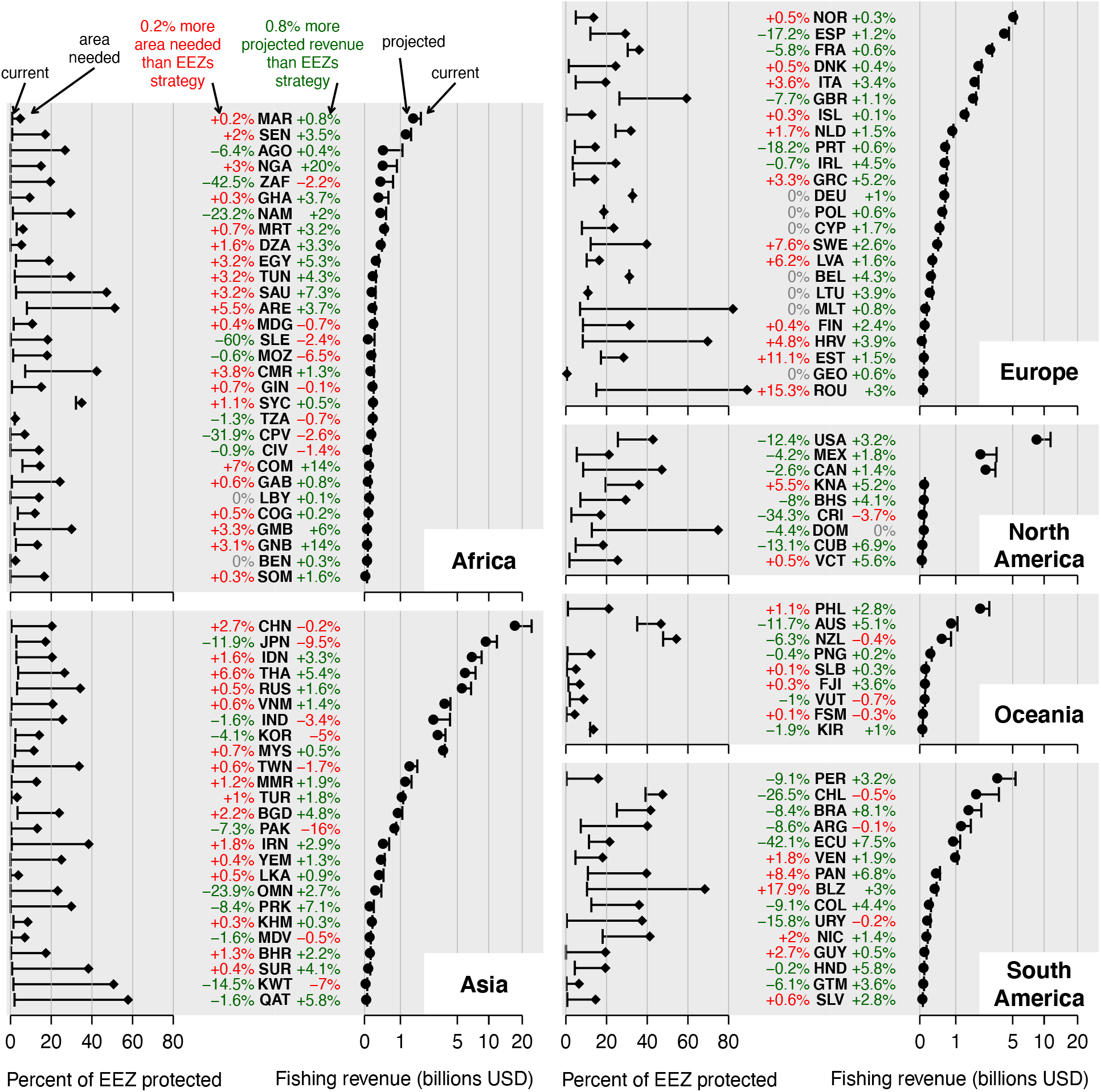
Current levels and required changes in protected area and expected fishing revenue necessary for each country to meet conservation targets, under a *MinCost* objective and all-oceans strategy. Percentage labels assess savings (green: absolute % difference) and cost (red) of the global-ocean strategy compared to the EEZs strategy. Countries are ordered by decreasing size of total fishing revenue (countries with revenue less than 50 million USD not shown).

A global-ocean strategy offered more pronounced benefits regarding the impacts of proposed MPAs on national fisheries revenues from domestic and distant water fleets, with 73% of countries seeing lower opportunity costs of foregoing catch. The relative difference in revenue retained under the global-ocean compared to EEZs strategy is comparable to areal saving (mean = 2.1%, SD = 5.1%), and would nonetheless currently amount to 4.75B USD in annual global savings. Notable economic beneficiaries from a global-ocean strategy include Nigeria (NGA), Comoros (COM), and Guinea-Bissau (GSB). Some 20% of countries are poised to incur lower revenues, with Pakistan (PAK), Japan (JPN), and Kuwait (KWT) most affected, spurred by loss of revenue within their EEZs as well as potential decreases in fishing opportunities for distant water fleets. For the majority of countries, fishing revenue savings suggest there is a large economic incentive toward global coordination and cooperation of MPA planning (*30*).

The MPA commitments necessary for sufficient biodiversity representation under the global-ocean strategy differed widely among countries. There was no correlation between additional MPA requirements and the amount of current protection (Spearman’s ρ = 0.02), but substantial additions are needed for most countries, with some requiring more than 50% of EEZ area (e.g. Romania (ROU), Malta (MLT), Dominican Republic (DOM)), especially when minimizing fisheries revenue loss (Fig. 4). In the mechanisms championing a 30% by 2030 MPA target in support of biodiversity, the recognition of the different national contributions required to address the overall shared responsibility is key. The appreciation that larger areal percentages, when carefully placed, will almost always be the economically prudent choice is equally important.

The total global cost of even the most economical approach to safeguarding marine biodiversity (global-oceans *MinCost* strategy) is substantial, at roughly 29.5 billion USD, or 18.7% in current annual revenue forgone. As with area, this cost varies strongly across nations (mean = −24.1%, SD = 21.8%). These revenue losses, however, are potentially mitigated by the known increase in fisheries productivity MPAs have from direct spill-over effects and refugia function for depleted, vulnerable, and overexploited stocks (*31, 32*). There are also many additional potential benefits from MPAs that are not included in this study, e.g., climate mitigation and adaptation (*22, 33*). In many countries fishing comprises a relatively small proportion of overall GDP, but for low income and emerging market economies, even small impacts to fisheries may pose threats to food security and livelihoods, especially in coastal countries where people are heavily dependent on small-scale fisheries for their nutritional supply and local economy (*34*). Our findings highlight the need for sustainable financing or other coordinated financial cooperation to assist countries in their contribution to global marine biodiversity conservation efforts (*16, 35*).

For the high seas, full agreement on an MPA network has remained elusive due to the required regulation of national industries operating in extra-jurisdictional MPAs and limited clarity on conservation outcomes (*15, 36–38*). There have been calls to close the high seas to fishing altogether, given their importance as reservoirs for high value species (*39*). However, our results show that, if in lieu of increasing protection within EEZs, a conservation focus on just the high seas would leave a large portion of biodiversity insufficiently safeguarded, including species restricted to EEZs and those cross-jurisdictional with the high seas (Fig. 3 middle). Full closure of the high seas therefore cannot substitute for the need for MPAs within EEZs. Importantly, this study suggests that strategic protection of select high seas areas while leaving others open to existing fishing activities will facilitate coordinated global conservation efforts by providing a substantial economic buffer to countries and easing the conservation burden within their EEZs.

Our study offers a roadmap for global ocean conservation. But, as other work, it retains assumptions regarding existing MPA effectiveness, species-level conservation targets, and very fine-scale occurrences that will benefit from future updates and extensions. The addition of further taxa with currently too limited and biased taxonomic representation will expand the recommended MPA network and likely change the among-nation distribution. Our economic assessment uses the most authoritative and spatially refined global catch data, but its reliability does vary among sectors and governments (*25*) and fish prices used in the analyses may be biased toward industrial rather than artisanal fisheries (*24, 26*). This study can also serve as a foundation to incorporate additional social and cultural dimensions of dependent human communities in assessing the costs and benefits of different MPAs settings; such information would be important in integrating social equity in discussing a globally-coordinated MPA network.

Our findings underscore the urgency and pay-off of global coordination to ensure the persistence of marine biodiversity. This is a pivotal time to re-evaluate ocean governance frameworks and position fisheries economies on a progressive path towards sustainability. There is an urgent need for nations to embrace their MPA estate and fishing economies as part of a complementary global network of the commons. Dedicated engagement from UNCLOS and mechanisms associated with the post-2020 global biodiversity framework can provide vital support for nations globally to benefit from a shared approach. Our framework pinpoints critical protection priorities, their effects on fisheries revenues and rationale, and highlights the substantial gains possible from a collaborative and quantitatively guided approach to marine biodiversity conservation.

## Materials and Methods

### Marine Protected Areas (MPAs)

We used the World Database on Protected Areas (December 2020 WDPA monthly release) to delineate MPAs (*11*). We excluded PAs that did not have designated, inscribed, or established status, points without a reported area, terrestrial reserves, and UNESCO Man and Biosphere Reserves. The remaining points were buffered to a size equal to the reported area (*41*), combined with the polygons and dissolved to create a global MPA layer. This layer was then rasterized to the same resolution and extent as the human modification data, and used to remove any HM 1-km pixels located inside MPAs so that currently protected areas did not contribute to aggregated summaries of HM. We acknowledge that coverage alone incompletely captures actual conservation progress as MPA objectives, resourcing and management varies widely from country to country (*42, 43*). Our framework can be updated as alternative databases become sufficiently comprehensive for both MPA effectiveness and Other Area-based Conservation Measures (OECMs).

### Human impacts

We used thirteen layers of abatable human impacts from Halpern *et al*. (*2*) to calculate a global map of marine cumulative abatable human modification. We used bilinear interpolation to align the extents of the layers and reprojected the layers to a 1km-pixel Behrmann equal-area projection. We then log(*x* + 1)-transformed and normalized the layers to a 0-1 range. We calculated cumulative human modification HM using a fuzzy algebraic sum, given by

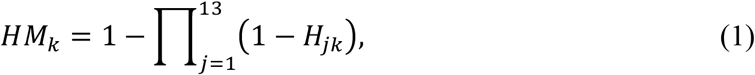

where *H_jk_* represents the log-normalized value of human impact layer *j* at location *k*. The fuzzy sum makes the contribution of a given stressor decrease as values from other stressors co-occur (*44*). *HM_k_* will be at least as large as the largest individual stressor *H_jk_*, but the additional contributions of other variables decrease the more they overlap. From this, we calculated the mean HM value by cell in a global equal-area grid (approximately 55×55 km at 30° latitudes).

### Species data

We used predicted range maps of 12,927 species of marine fishes and 119 species of marine mammals available through AquaMaps (www.aquamaps.org) (*45*). This comprises the most taxonomically comprehensive representation (>75% extant diversity) available for both these groups and for any larger marine species group. Other widely distributed marine taxa with more than several hundred species currently remain insufficiently mapped (<50% species coverage) to include without introducing significant spatial biases. Range maps provided probabilities of occurrence within a global 0.5°x0.5° grid, based on a combination of environmental niche modeling and expert review. These maps were projected to the Behrmann equal-area grid using bilinear interpolation, and then interpreted as binary presence/absence maps with a probability of occurrence threshold of 0.5 (representing species’ core ranges). Closing taxonomic and spatial knowledge gaps is vital for comprehensive biodiversity conservation (*46*), and further work is needed to expand our analyses to other species groups, but evidence of the close concordance between marine vertebrate and invertebrate biodiversity patterns (*45*) suggests that future inclusion of other taxa will yield qualitatively similar results.

### Fisheries revenue

The *Sea Around Us* (SAU) (www.seaaroundus.org) project’s reconstructed catch database aims to provide full accounting of fisheries catches by including catches from subsistence and recreational fishing sectors, discards and when possible, illegal, unreported and unregulated (IUU) fishing, which, by definition, are not part of official national data reported to the Food and Agriculture Organization of the United Nations (FAO) (*25*). The SAU classifies catch data by sector: industrial, artisanal, subsistence and recreational and provides time series catch data (1950 – 2015). Data are provided at half degree grid cells and each catch record is associated with taxon, fishing country and year (*24*).

Ex-vessel prices are the prices that fishers receive directly for their catch, or the selling price when it first enters the supply chain. Ex-vessel price data of each taxon by year and fishing country was extracted from the Fisheries Economic Research Unit (FERU) global ex-vessel price database (*26, 47, 48*). The detailed method for building the global ex-vessel database can be found in Tai et al., 2017. The updated price was matched with each record of the catch data in the SAU reconstructed catch database (*25*). By combining the catch data with the fishing ex-vessel price data of each marine taxon, the landed values can be estimated for different fishing countries at different spatial locations.

### Optimization

We sought to identify global reserve networks that meet targets set for the core ranges of P = 13,046 species of marine fish and mammals. We modeled the most efficient trajectories to optimality for each scenario by starting with the current MPA network and successively adding areas by greatest total contribution to conservation targets. We used a linear program (LP) problem definition with N = 121,722 decision variables representing cells in the global 55×55 km equal-area grid containing marine habitat in the form:

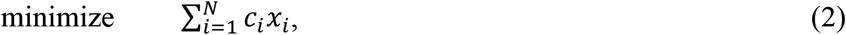

subject to:

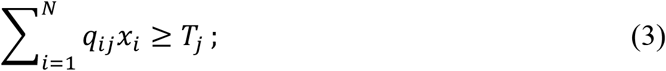

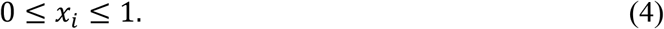

Equation (2) minimizes the cost of the reserve network. We defined two different cost variables. The first minimized reserve network area (*MinArea* scenario); the second minimized the fishing opportunity cost (*MinCost* scenario). The decision variables *x_i_* determine the proportion of habitat in each cell *i* that is included in the reserve network. Cell costs were calculated as

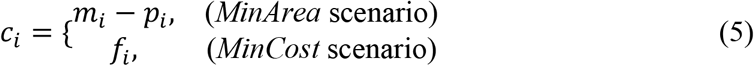

where *m_i_* represents the proportion of marine area in cell *i* and *p_i_* is the proportion of area of cell *i* that is currently protected.

The inequalities represented in Eq. (3) ensure that the amount of area protected by the reserve network is equal to or above a specified threshold *T_j_* of the total range size for each species *j*. The amount of habitat *q_ij_* “available” for future protection for species *j* in cell *i* was calculated as

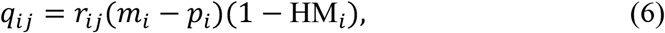

where *r_ij_* is a binary variable indicating whether or not cell *i* overlaps with the range of species *j*, and HM_*i*_ is the mean HM value in cell *i*. The conservation threshold *T_j_* was calculated as a function of a species’ total range size with a piecewise log-linear function that specified representation targets of 100% for species with the smallest ranges and 10% for species with the largest ranges (*49, 50*), and is given by

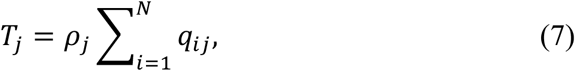

with *ρ_j_* the proportion of range area determined by the species representation target, following a commonly used target setting approach for terrestrial species (*49*). As global marine area is much larger than terrestrial area and marine species generally occupy larger areas, we used larger area of habitat thresholds to categorize rare (<100,000 km^2^) and widespread (>2,500,000 km^2^) species.

## Supporting information

Supplemental Data 1 - Study species

Supplemental Data 2 - Countries

## Acknowledgments

The authors would like to thank John Wilshire and Kate Ingenloff of Map of Life for help with data curation, and Greta Carrete Vega, Tamara Huete, and Javier Alba of Vizzuality for assistance with data visualization.

## Funding

This work was supported by the E.O. Wilson Biodiversity Foundation in association with the Half-Earth Project. W.J. acknowledges support from NSF grant DEB-1441737, and NASA grants 80NSSC17K0282 and 80NSSC18K0435.

## Author contributions

**D. Scott Rinnan:** Conceptualization, Formal analysis, Investigation, Methodology, Project administration, Software, Validation, Visualization, Writing – original draft. **Gabriel Reygondeau:** Conceptualization, Data curation, Formal analysis, Investigation, Methodology, Visualization, Writing – review & editing. **Jennifer McGowan:** Investigation, Writing – review & editing. **Vicky Lam:** Data curation, Formal analysis, Resources, Writing – review & editing. **Rashid Sumaila:** Data curation, Resources, Writing – review & editing. **Ajay Ranipeta:** Data curation, Resources. **Kristin Kaschner:** Data curation, Resources, Writing – review & editing. **Cristina Garilao:** Data curation, Resources. **William L. Cheung:** Supervision, Writing – review & editing. **Walter Jetz:** Conceptualization, Funding acquisition, Project administration, Supervision, Visualization, Writing – review & editing.

## Competing interests

Authors declare no competing interests.

## Data and materials availability

Species distribution data is available from the AquaMaps database via www.aquamaps.org. Fishing value data is available from the Sea Around Us database via www.seaaroundus.org. Human impacts data is available via doi:10.5063/F11Z42N8. Prioritization results by species and country and species representation targets are available as supplementary materials.

---

**Table S1.** Human impact layers used to calculate cumulative human modification.

| Commercial fishing categories | Ocean-based drivers | Land-based drivers |
| --- | --- | --- |
| Demersal destructive fishing | Artisanal fishing | Plumes (fertilizer) |
| Demersal nondestructive high bycatch | Species invasion | Plumes (pesticides) |
| Demersal nondestructive low bycatch | Oil rigs | Nonpoint inorganic pollution |
| Pelagic high bycatch |  | Nonpoint organic pollution |
| Pelagic low bycatch |  | Population |

**Fig. S1.**
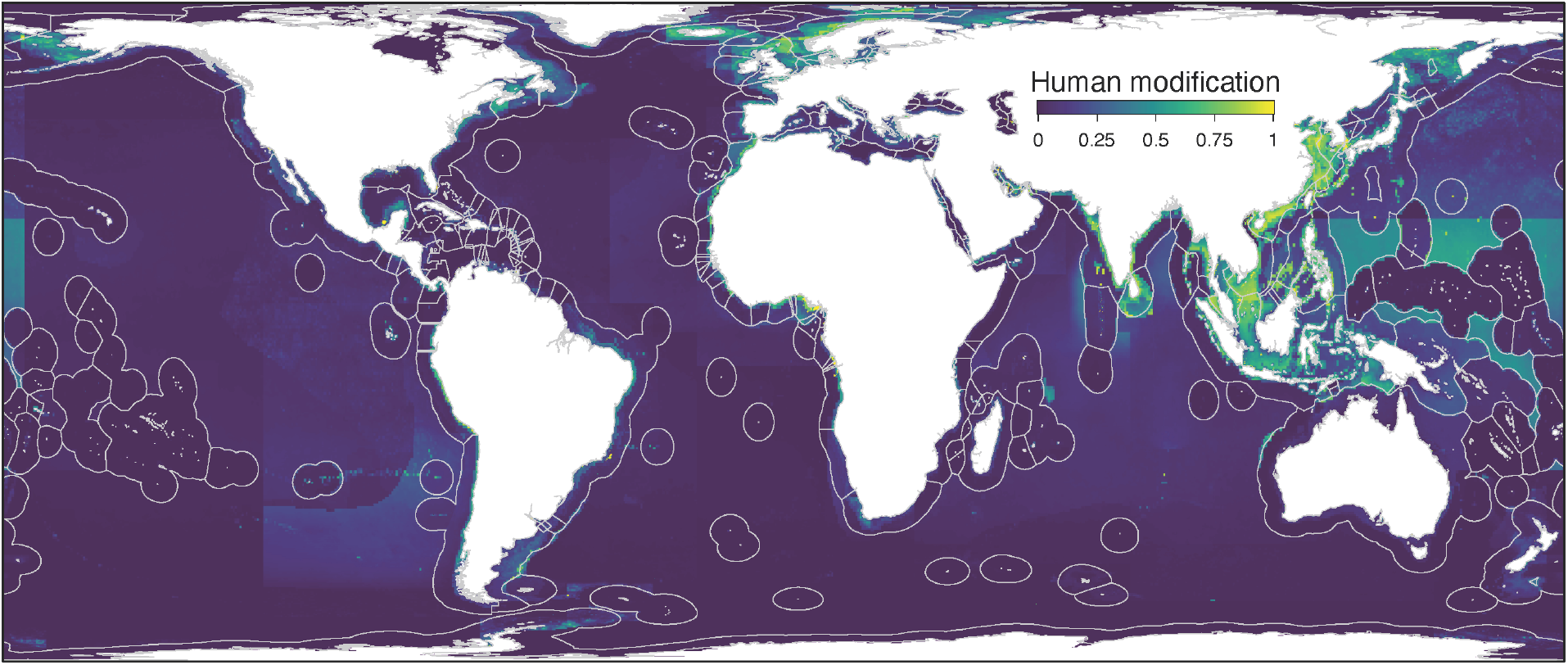
Cumulative abatable human modification of the global ocean, summarizing thirteen different types of marine human impacts.

**Fig. S2.**
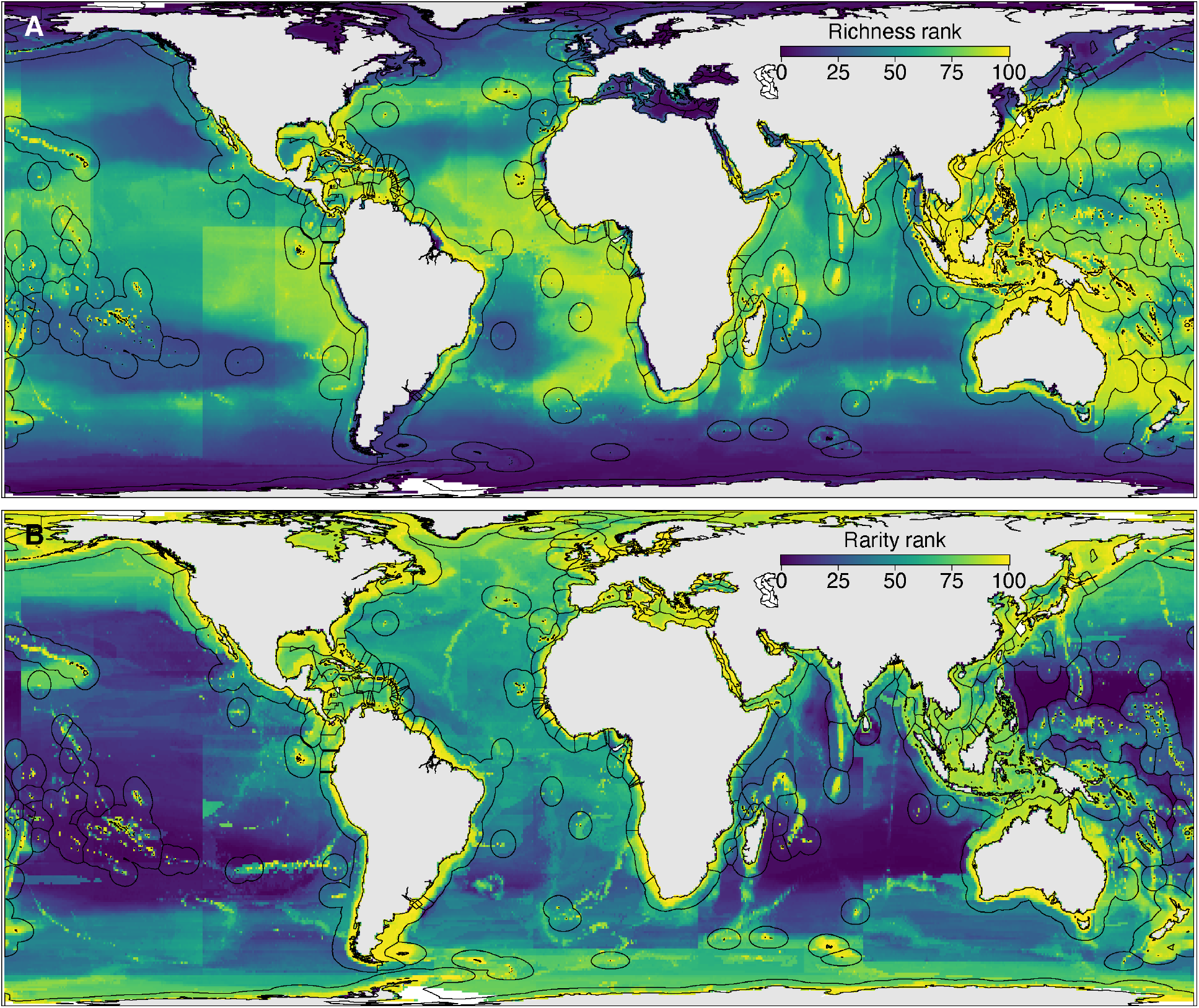
Patterns of species (A) richness and (B) range size rarity of 13,046 marine vertebrates, comprising 12,927 fishes and 119 mammals.

**Fig. S3.**
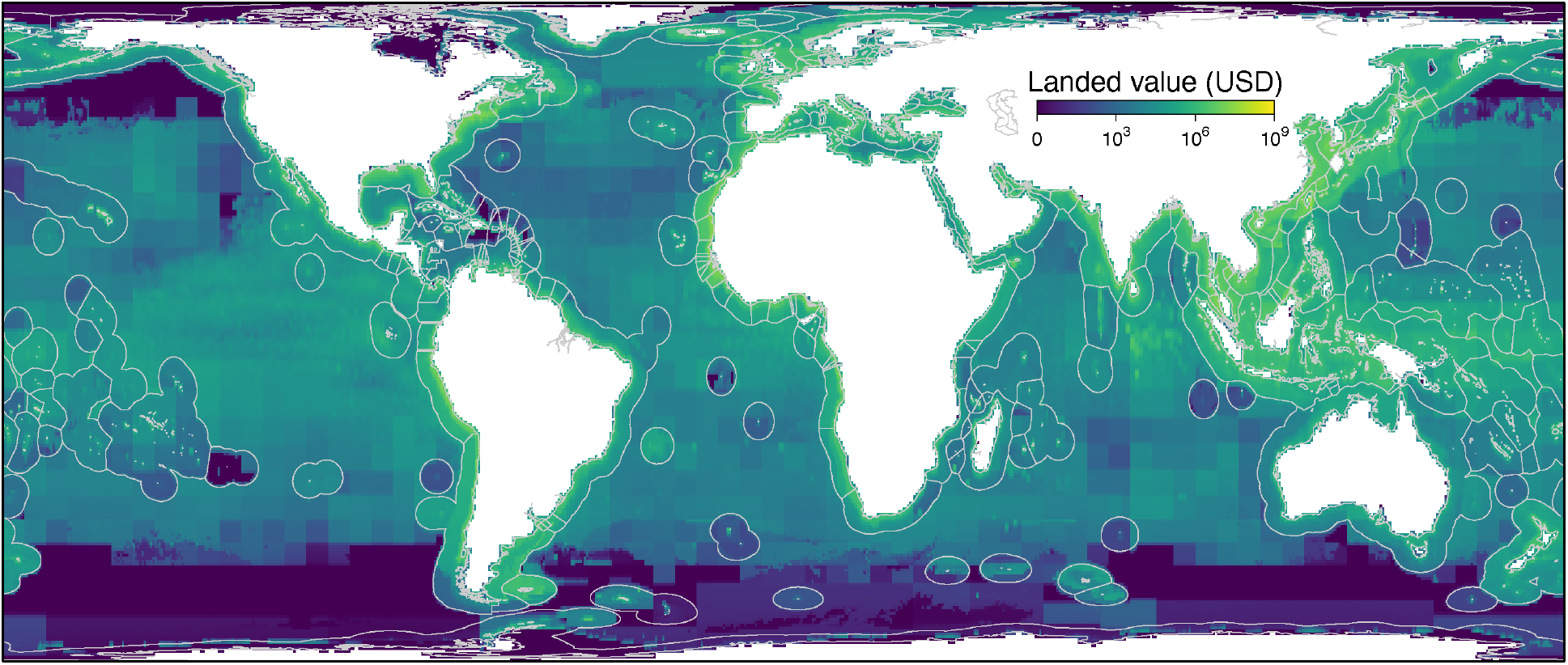
Landed value estimates in real 2010 USD, averaged from 2007–2016 and aggregated across fisheries.

## Footnotes

1 Real 2010 USD value of total global landed value averaged 2007-2016 and adjusted to 2021.

## Notes

### Competing Interest Statement

The authors have declared no competing interest.

## References and Notes

1. B. S. Halpern, S. Walbridge, K. A. Selkoe, C. V. Kappel, F. Micheli, C. D’Agrosa, J. F. Bruno, K. S. Casey, C. Ebert, H. E. Fox, R. Fujita, D. Heinemann, H. S. Lenihan, E. M. P. Madin, M. T. Perry, E. R. Selig, M. Spalding, R. Steneck, R. Watson, A Global Map of Human Impact on Marine Ecosystems. Science. 319, 948–952 (2008).

2. B. S. Halpern, M. Frazier, J. Potapenko, K. S. Casey, K. Koenig, C. Longo, J. S. Lowndes, R. C. Rockwood, E. R. Selig, K. A. Selkoe, S. Walbridge, Spatial and temporal changes in cumulative human impacts on the world’s ocean. Nat Commun. 6, 7615 (2015).

3. N. L. Bindoff, W. W. Cheung, J. G. Kairo, J. Arstegui, V. A. Guinder, R. Hallberg, N. Hilmi, N. Jiao, M. S. Karim, L. Levin, in IPCC special report on the ocean and cryosphere in a changing climate (2019).

4. S. M. Díaz, J. Settele, E. Brondízio, H. Ngo, M. Guèze, J. Agard, A. Arneth, P. Balvanera, K. Brauman, S. Butchart, K. M. A. Chan, L. A. Garibaldi, K. Ichii, J. Liu, S. Subramanian, G. Midgley, P. Miloslavich, Z. Molnár, D. Obura, A. Pfaff, S. Polasky, A. Purvis, J. Razzaque, B. Reyers, R. Roy Chowdhury, Y.-J. Shin, I. Visseren-Hamakers, K. Willis, C. Zayas, The global assessment report on biodiversity and ecosystem services: Summary for policy makers (Intergovernmental Science-Policy Platform on Biodiversity and Ecosystem Services, 2019; http://ri.conicet.gov.ar/handle/11336/116171).

5. D. Pauly, W. W. L. Cheung, Sound physiological knowledge and principles in modeling shrinking of fishes under climate change. Global Change Biology. 24, e15–e26 (2018).

6. E. Sala, N. Knowlton, Global Marine Biodiversity Trends. Annual Review of Environment and Resources. 31, 93–122 (2006).

7. D. J. McCauley, M. L. Pinsky, S. R. Palumbi, J. A. Estes, F. H. Joyce, R. R. Warner, Marine defaunation: Animal loss in the global ocean. Science. 347 (2015), doi:10.1126/science.1255641.

8. V. Christensen, M. Coll, C. Piroddi, J. Steenbeek, J. Buszowski, D. Pauly, A century of fish biomass decline in the ocean. Marine Ecology Progress Series. 512, 155–166 (2014).

9. U. R. Sumaila, T. C. Tai, V. W. Y. Lam, W. W. L. Cheung, M. Bailey, A. M. Cisneros-Montemayor, O. L. Chen, S. S. Gulati, Benefits of the Paris Agreement to ocean life, economies, and people. Science Advances. 5, eaau3855 (2019).

10. E. Sala, S. Giakoumi, No-take marine reserves are the most effective protected areas in the ocean. ICES J Mar Sci. 75, 1166–1168 (2018).

11. UNEP-WCMC and IUCN, Protected Planet: The World Database on Protected Areas (WDPA). Protected Planet: The World Database on Protected Areas (WDPA) (2019), (available at https://www.protectedplanet.net).

12. C. J. Klein, C. J. Brown, B. S. Halpern, D. B. Segan, J. McGowan, M. Beger, J. E. M. Watson, Shortfalls in the global protected area network at representing marine biodiversity. Scientific Reports. 5, 17539 (2015).

13. K. Jantke, K. R. Jones, J. R. Allan, A. L. Chauvenet, J. E. Watson, H. P. Possingham, Poor ecological representation by an expensive reserve system: Evaluating 35 years of marine protected area expansion. Conservation Letters. 11, e12584 (2018).

14. E. O. Wilson, Half-earth: our planet’s fight for life (WW Norton & Company, 2016).

15. B. C. O’Leary, H. L. Allen, K. L. Yates, R. W. Page, A. W. Tudhope, C. McClean, C. M. Roberts, 30X30 A Blueprint for Ocean Protection. How we can protect. 30 (2019).

16. A. Waldron, V. Adams, J. Allan, A. Arnell, G. Asner, S. Atkinson, A. Baccini, J. Baillie, A. Balmford, J. Austin Beau, Protecting 30% of the planet for nature: costs, benefits and economic implications (2020).

17. UNEP, Convention on Biological Diversity: Update of the zero draft of the post-2020 global biodiversity framework (2020).

18. K. R. Jones, C. J. Klein, B. S. Halpern, O. Venter, H. Grantham, C. D. Kuempel, N. Shumway, A. M. Friedlander, H. P. Possingham, J. E. M. Watson, The Location and Protection Status of Earth’s Diminishing Marine Wilderness. Current Biology. 28, 2506–2512.e3 (2018).

19. K. R. Jones, C. J. Klein, H. S. Grantham, H. P. Possingham, B. S. Halpern, N. D. Burgess, S. H. M. Butchart, J. G. Robinson, N. Kingston, N. Bhola, J. E. M. Watson, Area Requirements to Safeguard Earth’s Marine Species. One Earth. 2, 188–196 (2020).

20. M. E. Visalli, B. D. Best, R. B. Cabral, W. W. L. Cheung, N. A. Clark, C. Garilao, K. Kaschner, K. Kesner-Reyes, V. W. Y. Lam, S. M. Maxwell, J. Mayorga, H. V. Moeller, L. Morgan, G. O. Crespo, M. L. Pinsky, T. D. White, D. J. McCauley, Data-driven approach for highlighting priority areas for protection in marine areas beyond national jurisdiction. Marine Policy, 103927 (2020).

21. Q. Zhao, F. Stephenson, C. Lundquist, K. Kaschner, D. Jayathilake, M. J. Costello, Where Marine Protected Areas would best represent 30% of ocean biodiversity. Biological Conservation. 244, 108536 (2020).

22. E. Sala, J. Mayorga, D. Bradley, R. B. Cabral, T. B. Atwood, A. Auber, W. Cheung, C. Costello, F. Ferretti, A. M. Friedlander, S. D. Gaines, C. Garilao, W. Goodell, B. S. Halpern, A. Hinson, K. Kaschner, K. Kesner-Reyes, F. Leprieur, J. McGowan, L. E. Morgan, D. Mouillot, J. Palacios-Abrantes, H. P. Possingham, K. D. Rechberger, B. Worm, J. Lubchenco, Protecting the global ocean for biodiversity, food and climate. Nature, 1–6 (2021).

23. FAO, The State of World Fisheries and Aquaculture 2020: Sustainability in action (FAO, Rome, Italy, 2020; http://www.fao.org/documents/card/en/c/ca9229en), The State of World Fisheries and Aquaculture (SOFIA).

24. D. Zeller, M. L. D. Palomares, A. Tavakolie, M. Ang, D. Belhabib, W. W. L. Cheung, V. W. Y. Lam, E. Sy, G. Tsui, K. Zylich, D. Pauly, Still catching attention: Sea Around Us reconstructed global catch data, their spatial expression and public accessibility. Marine Policy. 70, 145–152 (2016).

25. D. Pauly, D. Zeller, Catch reconstructions reveal that global marine fisheries catches are higher than reported and declining. Nature Communications. 7, 1–9 (2016).

26. T. C. Tai, T. Cashion, V. W. Y. Lam, W. Swartz, U. R. Sumaila, Ex-vessel Fish Price Database: Disaggregating Prices for Low-Priced Species from Reduction Fisheries. Front. Mar. Sci. 4 (2017), doi:10.3389/fmars.2017.00363.

27. D. C. Dunn, C. Jablonicky, G. O. Crespo, D. J. McCauley, D. A. Kroodsma, K. Boerder, K. M. Gjerde, P. N. Halpin, Empowering high seas governance with satellite vessel tracking data. Fish and Fisheries. 19, 729–739 (2018).

28. M. Pinsky, G. Reygondeau, R. Caddell, J. Palacios Abrantes, J. Spijkers, W. Cheung, Preparing ocean governance for species on the move. Science. 360 (2018), doi:10.1126/science.aat2360.

29. J. Palacios-Abrantes, G. Reygondeau, C. C. C. Wabnitz, W. W. L. Cheung, The transboundary nature of the world’s exploited marine species. Scientific Reports. 10, 17668 (2020).

30. K. A. Miller, G. R. Munro, U. R. Sumaila, W. W. L. Cheung, Governing Marine Fisheries in a Changing Climate: A Game-Theoretic Perspective. Canadian Journal of Agricultural Economics/Revue canadienne d’agroeconomie. 61, 309–334 (2013).

31. B. S. Halpern, S. E. Lester, J. B. Kellner, Spillover from marine reserves and the replenishment of fished stocks. Environmental Conservation. 36, 268–276 (2009).

32. M. D. Lorenzo, P. Guidetti, A. D. Franco, A. Calò, J. Claudet, Assessing spillover from marine protected areas and its drivers: A meta-analytical approach. Fish and Fisheries. 21, 906–915 (2020).

33. C. M. Roberts, B. C. O’Leary, D. J. McCauley, P. M. Cury, C. M. Duarte, J. Lubchenco, D. Pauly, A. Sáenz-Arroyo, U. R. Sumaila, R. W. Wilson, B. Worm, J. C. Castilla, Marine reserves can mitigate and promote adaptation to climate change. PNAS. 114, 6167–6175 (2017).

34. C. D. Golden, E. H. Allison, W. W. L. Cheung, M. M. Dey, B. S. Halpern, D. J. McCauley, M. Smith, B. Vaitla, D. Zeller, S. S. Myers, Nutrition: Fall in fish catch threatens human health. Nature News. 534, 317 (2016).

35. J. McGowan, R. Weary, L. Carriere, E. T. Game, J. L. Smith, M. Garvey, H. P. Possingham, Prioritizing debt conversion opportunities for marine conservation. Conservation Biology. 34, 1065–1075 (2020).

36. B. C. O’Leary, R. L. Brown, D. E. Johnson, H. von Nordheim, J. Ardron, T. Packeiser, C. M. Roberts, The first network of marine protected areas (MPAs) in the high seas: The process, the challenges and where next. Marine Policy. 36, 598–605 (2012).

37. E. M. De Santo, Á. Ásgeirsdóttir, A. Barros-Platiau, F. Biermann, J. Dryzek, L. R. Gonçalves, R. E. Kim, E. Mendenhall, R. Mitchell, E. Nyman, M. Scobie, K. Sun, R. Tiller, D. G. Webster, O. Young, Protecting biodiversity in areas beyond national jurisdiction: An earth system governance perspective. Earth System Governance. 2, 100029 (2019).

38. R. Tiller, E. De Santo, E. Mendenhall, E. Nyman, The once and future treaty: Towards a new regime for biodiversity in areas beyond national jurisdiction. Marine Policy. 99, 239–242 (2019).

39. U. R. Sumaila, V. W. Y. Lam, D. D. Miller, L. Teh, R. A. Watson, D. Zeller, W. W. L. Cheung, I. M. Côté, A. D. Rogers, C. Roberts, E. Sala, D. Pauly, Winners and losers in a world where the high seas is closed to fishing. Scientific Reports. 5, 8481 (2015).

40. UNEP-WCMC, IUCN and NGS, “Protected Planet Report 2018” (UNEP-WCMC, IUCN and NGS: Cambridge UK; Gland, Switzerland; and Washington, D.C., USA., 2018).

41. P. Visconti, M. Di Marco, J. G. Álvarez-Romero, S. R. Januchowski-Hartley, R. L. Pressey, R. Weeks, C. Rondinini, Effects of Errors and Gaps in Spatial Data Sets on Assessment of Conservation Progress: Errors and Gaps in Spatial Data Sets. Conserv. Biol. 27, 1000–1010 (2013).

42. D. A. Gill, M. B. Mascia, G. N. Ahmadia, L. Glew, S. E. Lester, M. Barnes, I. Craigie, E. S. Darling, C. M. Free, J. Geldmann, S. Holst, O. P. Jensen, A. T. White, X. Basurto, L. Coad, R. D. Gates, G. Guannel, P. J. Mumby, H. Thomas, S. Whitmee, S. Woodley, H. E. Fox, Capacity shortfalls hinder the performance of marine protected areas globally. Nature. 543, 665–669 (2017).

43. M. D. Barnes, L. Glew, C. Wyborn, I. D. Craigie, Prevent perverse outcomes from global protected area policy. Nature Ecology & Evolution. 2, 759–762 (2018).

44. C. M. Kennedy, J. R. Oakleaf, D. M. Theobald, S. Baruch-Mordo, J. Kiesecker, Managing the middle: A shift in conservation priorities based on the global human modification gradient. Glob Change Biol. 25, 811–826 (2019).

45. K. Kaschner, K. Kesner-Reyes, C. Garilao, J. Rius-Barile, T. Rees, R. Froese, AquaMaps: Predicted range maps for aquatic species (2019), (available at https://www.aquamaps.org).

46. M. J. Costello, C. Chaudhary, Marine Biodiversity, Biogeography, Deep-Sea Gradients, and Conservation. Current Biology. 27, R511–R527 (2017).

47. U. R. Sumaila, A. D. Marsden, R. Watson, D. Pauly, A Global Ex-vessel Fish Price Database: Construction and Applications. J Bioecon. 9, 39–51 (2007).

48. W. Swartz, R. Sumaila, R. Watson, Global Ex-vessel Fish Price Database Revisited: A New Approach for Estimating ‘Missing’ Prices. Environ Resource Econ. 56, 467–480 (2013).

49. A. S. L. Rodrigues, H. R. Akçakaya, S. J. Andelman, M. I. Bakarr, L. Boitani, T. M. Brooks, J. S. Chanson, L. D. C. Fishpool, G. A. B. Da Fonseca, K. J. Gaston, M. Hoffmann, P. A. Marquet, J. D. Pilgrim, R. L. Pressey, J. Schipper, W. Sechrest, S. N. Stuart, L. G. Underhill, R. W. Waller, M. E. J. Watts, X. Yan, Global Gap Analysis: Priority Regions for Expanding the Global Protected-Area Network. BioScience. 54, 1092 (2004).

50. D. S. Rinnan, W. Jetz, Terrestrial conservation opportunities and inequities revealed by global multi-scale prioritization. bioRxiv, in press, doi:10.1101/2020.02.05.936047.

